# Mitochondrial nicotinic acetylcholine receptors form complexes with Bax upon apoptosis induction

**DOI:** 10.1101/2020.05.13.094102

**Authors:** Olena Kalashnyk, Olena Lykhmus, Kateryna Uspenska, Mykhailo Izmailov, Sergiy Komisarenko, Maryna Skok

**Author notes:** Corresponding author (MS).

## Abstract

Nicotinic acetylcholine receptors (nAChRs) mediate fast synaptic transmission in muscles and autonomic ganglia and regulate cytokine and neurotransmitter release in the brain and nonexcitable cells. The nAChRs expressed in the outer membrane of mitochondria control the early events of mitochondria-driven apoptosis like cytochrome c release by affecting intramitochondrial kinase pathways. However, the mechanisms through which nAChRs influence mitochondrial permeability remain obscure. Previously we demonstrated that mitochondrial nAChRs interact with voltage-dependent anion channels (VDAC) involved in forming the pore in mitochondria membrane. Here we put an aim to explore the connection of nAChRs to pro-apoptotic protein Bax and its changes in the course of apoptosis induction. By using molecular modeling in silico, it was shown that both Bax and VDAC bind within the 4^th^ transmembrane portion of nAChR subunits. Experimentally, α7 nAChR-Bax and α7 nAChR-VDAC complexes were identified by sandwich ELISA in mitochondria isolated from astrocytoma U373 cells. Stimulating apoptosis of U373 cells by 1μM H_2_O_2_ disrupted α7-VDAC complexes and favored formation of α7-Bax complexes. α7-selective agonist PNU282987 and type 2 positive allosteric modulator PNU120596 disrupted α7-Bax and returned α7 nAChR to complex with VDAC. It is concluded that mitochondrial nAChRs regulate apoptosis-induced mitochondrial channel formation by modulating the interplay of apoptosis-related proteins in mitochondria outer membrane.

## Introduction

Nicotinic acetylcholine receptors (nAChRs) were initially discovered in muscles and autonomic ganglia where they function as ligand-gated ion channels mediating fast synaptic transmission[1–2]. Further they were found in the brain [3] and in many non-excitable cells where they can function both iono- and metabotropically [4] to regulate vital cellular properties like survival, proliferation, secretion of neurotransmitters and cytokines [5–7]. Finally, the nAChRs were discovered in mitochondria to regulate the early events of mitochondria-driven apoptosis like cytochrome c (cyt c) release by affecting intramitochondrial kinases [8,9]. However, the mechanisms through which nAChRs influence mitochondrial permeability remain obscure. Previously we reported the connection of mitochondrial nAChRs to voltage-dependent anion channels (VDAC) [8] suggested to take part in regulating mitochondrial pore formation [10]. However, this role of VDAC was put under a doubt by the use of VDAC-knockout mice [11].

The alternative model suggests that mitochondrial pore releasing pro-apoptotic factors like cyt c, OMI/HTRA2, AC/DIABLO, and endonuclease G [12] is formed by pro-apoptotic Bcl-2 family proteins Bax or Bak, which oligomerize in the mitochondrial outer membrane. They are activated by the BH3 proteins that sense cellular stress and are inhibited by anti-apoptotic family members like Bcl-2 or Bcl-XL. As mentioned by Shamas-Din et al. [13], the interplay between members of the Bcl-2 family is determined by competing equilibria and largely depends on the conformational changes occurring due to specific conditions of cell physiology and interaction with other membrane proteins. Taking into account the established anti-apoptotic role of mitochondrial nAChRs, we suggested that they can participate in these interactions and put an aim to explore the connection of nAChRs to Bax in the course of apoptosis induction. Here we show that inducing apoptosis with H_2_O_2_ in astrocytoma U373 cells results in displacement of mitochondrial nAChRs from complexes with VDAC and formation of nAChR-Bax complexes followed by cyt c release. Treatment of mitochondria with α7 nAChR agonist PNU282987 disrupts nAChR-Bax complexes and restores nAChR-VDAC complexes that attenuates cyt c release. This data shed new light on the mechanisms of mitochondrial pore formation and its cholinergic regulation.

## Materials and methods

### *In silico* experiments

Interaction of nAChRs with Bax and VDAC was modeled using pyDockWEBweb server for the structural prediction of protein-protein interactions [14]. The protein structures were taken from RCSB Protein Data Base [15]:

nAChR PDB ID: 2BG9 (from *Torpedo marmorata)*
Bax PDB ID: 1F16 (from *Homo sapiens;* expressed in *Escherichia coli)*
VDAC PDB ID: 2JK4 (from *Homo sapiens;* expressed in *Escherichia coli)*

### Cells and reagents

All reagents and antibodies against VDAC and lamin were purchased from Sigma-Aldrich (Saint Louis, USA). Antibodies against α7(1-208) [16], α7(179-190) [17], and β2(190-200) [18] nAChR fragments and against cyt c [19] were obtained, validated and biotinylated previously in our lab. Neutravidin-peroxidase conjugate, Bax-specific, IRE-1α-specific antibodies, mitochondria isolation kit and IL-6 kit were from Invitrogen and were purchased from ALT Ukraine Ltd (representative of Thermo Fisher Scientific in Ukraine).

U373 cells (ATCC HTB-17) were from the stocks of Palladin Institute of Biochemistry. They were grown in 75 cm^2^ flasks at 37°C in RPMI1640 medium supplemented with 20 mM HEPES, 100 units/ml penicillin-streptomycin mixture and 10% FCS. Before the experiment, the cells were detached from the flask bottoms by gentle scratching, washed by centrifugation, counted and resuspended in the culture medium.

### Mitochondria isolation and experimental schedules

1. We used 10^7^ cells per one mitochondria portion. Mitochondria were isolated according to the instructions of the kit manufacturer. Their purity was characterized by ELISA using the antibodies against mitochondria-specific voltage-dependent anion channel (VDAC, [20]), nuclear-specific marker α-lamin B1 [21] and endoplasmic reticulum-specific marker IRE-1α [22] as described previously [23]. The isolated mitochondria were incubated with 0.9μM CaCl_2_ or 0.5 mM H_2_O_2_ for 5 min at RT, then pelleted by centrifugation at 7,000g, the supernatant being collected and tested in cyt c release ELISA test (see below).
2. Separate 10^7^ portions of U373 cells were incubated with 1mM H_2_O_2_ for 0.5, 1, 1.5 or 3 h, washed by centrifugation and mitochondria were isolated as above. The mitochondria pellet was frozen at −20°C, thawed and treated with lysing buffer (0.01 M Tris-HCl, pH 8.0; 0.14 NaCl; 0.025% NaN_3_; 1% Tween-20 and protease inhibitors cocktail) for 2 h on ice upon intensive stirring. The resulting lysates were pelleted by centrifugation (20 min at 20,000g). The protein concentration was established with the BCA Protein Assay kit. The resultant lysates were analysed by Sandwich ELISA for the presence of α7 nAChRs, α7-Bax complexes and cyt c as described below.
3. U373 cells were either pre-incubated or not with PNU282987 (1μM) or PNU120596 (10 μM) for 30 min, washed by centrifugation and then incubated with 1mM H_2_O_2_ for 1h. Mitochondria were isolated from both control and H_2_O_2_-treated cells and lysed as described above. Sandwich ELISA for the cyt c was performed in both mitochondria lysates and cell cytosole remained after mitochondria isolation.
4. U373 cells (4×10^7^) were incubated in the presence of 1mM H_2_O_2_ for 1h and the isolated mitochondria were divided into 4 portions. One portion remained intact, while three others were treated with 1% SDS, 30 nM PNU282987 or 300 nM PNU120596 for 15 min at RT, then washed by centrifugation and lysed as descried above. The resultant lysates were analyzed by ELISA for the levels of α7-Bax and β2-Bax complexes.
5. U373 cells were either incubated (2×10^7^) or not (1×10^7^) with 1mM H_2_O_2_ and mitochondria were isolated from both portions. Mitochondria of cells incubated with H_2_O_2_ were further divided into two equal portions and one portion was additionally treated with 30 nM PNU282987 for 15 min. Then mitochondria were frozen, lysed as described above and the resultant lysates were analyzed for the levels of α7 nAChRs, α7-Bax and α7-VDAC by Sandwich ELISA.

### Sandwich assays

To determine the level of α7nAChR subunits, the immunoplates (Nunc, MaxiSorp) were coated with rabbit α7(1-208)-specific antibody (20 μg/ml), blocked with 1% BSA, and the detergent lysates of the mitochondria were applied into the wells (1 μg of protein per 0.05 ml per well) for 2h at 37°C. The plates were washed with water and the second biotinylated α7(179-190)-specific antibody (6.6 μg/ml) was applied for additional 2 h at 37°C.

To evaluate the presence of the nAChR-Bax or nAChR-VDAC complexes, the plates were coated with α7(1-208)-specific antibody (20 μg/ml) and the bound Bax or VDAC were revealed with biotinylated Bax- (4 μg/ml) or VDAC-specific (3 μg/ml) antibodies.

To reveal α7-Bax or β2-Bax complexes, the plates were coated with α7(179-190)- or β2(190-200)-specific antibody (20 μg/ml) and the bound Bax was revealed with biotinylated Bax-specific antibody (4 μg/ml).

To measure cyt c released from mitochondria, the plates were coated with polyclonal cyt c-specific antibody (20 μg/ml) to catch the antigen from the lysed mitochondria (70 μg/ml),cytosole (150 μg/ml) or mitochondria supernatant (non-diluted). The bound cyt c was revealed with biotinylated cyt-c-specific antibody as described previously [8,19].

The bound biotinylated antibodies were revealed with Neutravidin-peroxidase conjugate and *o*-phenylendiamine-containing substrate solution. The optical density of ELISA plates was read at 490 nm using Stat-Fax 2000 ELISA Reader (Awareness Technologies, USA).

### The whole-cell assays

1. U373 cells were seeded into 96-well plates, 5×10^3^ per well, for attaching overnight. PNU282987 (1 μM) was added for 30 min, then the medium was replaced and 1 mM H_2_O_2_ was added for additional 1 h. For experiments with recombinant IL-6 1 mM H_2_O_2_ was added for 1h before IL-6 (100 pg/ml). After 24 h the supernatant was collected for IL-6 determination, while the cells were washed and tested in MTT assay [24].
2. The cells seeded as above were incubated in the presence of 1mM H_2_O_2_, 1μM PNU282987, their combination or LPS (1μg/ml) for 24 h. The supernatants were collected and analyzed for the presence of IL-6 using a commercial kit according to the manufacturer’s instructions.

### Statistical analysis

ELISA experiments have been performed in triplicates and mean values were used for statistical analysis using one-way ANOVA test and Origin 9.0 software. The data are presented as Mean±SD. Each experiment has been performed at least twice yielding similar results.

## Results

Top 10 models obtained using the pyDockWEB program demonstrated three potential binding sites of Bax and nAChR (Fig. 1A). All of them were located in the transmembrane portion of nAChR subunits, mostly in M4 domain (residues 450-460). Modeling nAChR-VDAC1 interaction resulted in a similar picture: top 10 models demonstrated potential binding of VDAC within M4 domain of nAChR (Fig. 1B).

**Fig 1.**
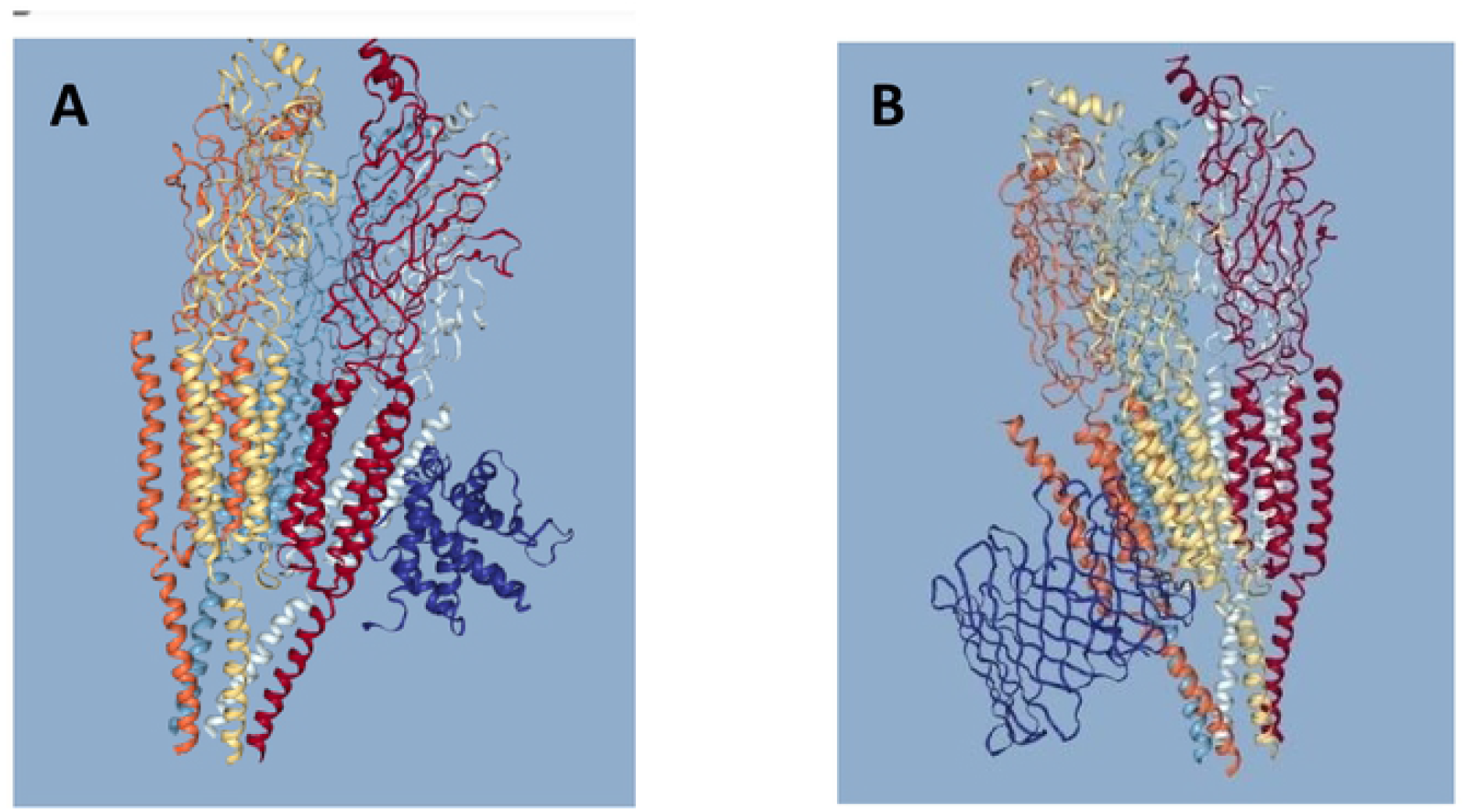
Molecular modeling of the nAChR-Bax (A) and nAChR-VDAC (B) interactions. Shown are models 1 from top 10 models calculated by pyDockWEB program.

To study if such complexes are really formed, we isolated mitochondria from U373 cells previously reported to contain several nAChR subtypes, including α7 ones [25]. Isolated mitochondria were VDAC-positive according to Sandwich ELISA (Fig. 2A) and released cyt c in response to treatment with either Ca^2+^ or H_2_O_2_ (Fig. 2B) as did mitochondria isolated from the mouse brain or liver [8, 26] and, therefore, were considered suitable for further functional studies.

**Fig 2.**
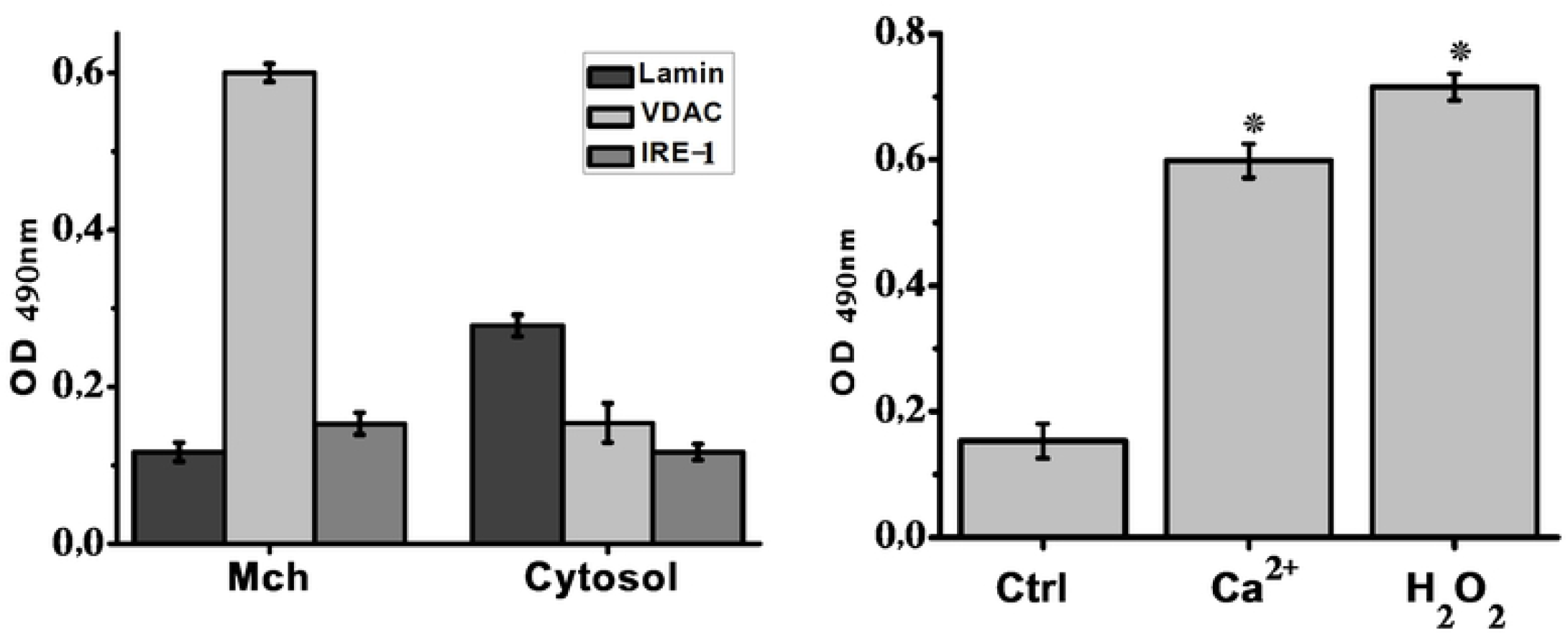
Characterization of mitochondria (Mch) isolated from U373 cells. Sandwich ELISA data for the expression of mitochondria-specific (VDAC), nucleus-specific (lamin) and endoplasmic reticulum-specific (IRE-1) markers (**A**) and cyt c release under the effect of 0.9 μM Ca^2+^ or 0.5 mM H_2_O_2_ (**B**). * – p<0.001 compared to non-treated mitochondria (Ctrl); n=3.

Mitochondria isolated from U373 cells cultured in the presence of 1 mM H_2_O_2_ during 0.5, 1 or 3h, were studied for the presence of α7 nAChRs, α7-Bax complexes and cyt c. As shown in Fig.3A, the level of α7 nAChR in mitochondria increased at 0.5h already and was maintained by 3h. The signal for α7-Bax increased at 1h and was also maintained during 2 hours more. The level of cyt c within mitochondria started to drop down at 0.5h and reached plateau by 1h(Fig.3B). Therefore, treatment of cells with H_2_O_2_ resulted in the increase of α7 nAChRs in mitochondria followed by formation of α7-Bax complexes and cyt c release.

**Fig 3.**
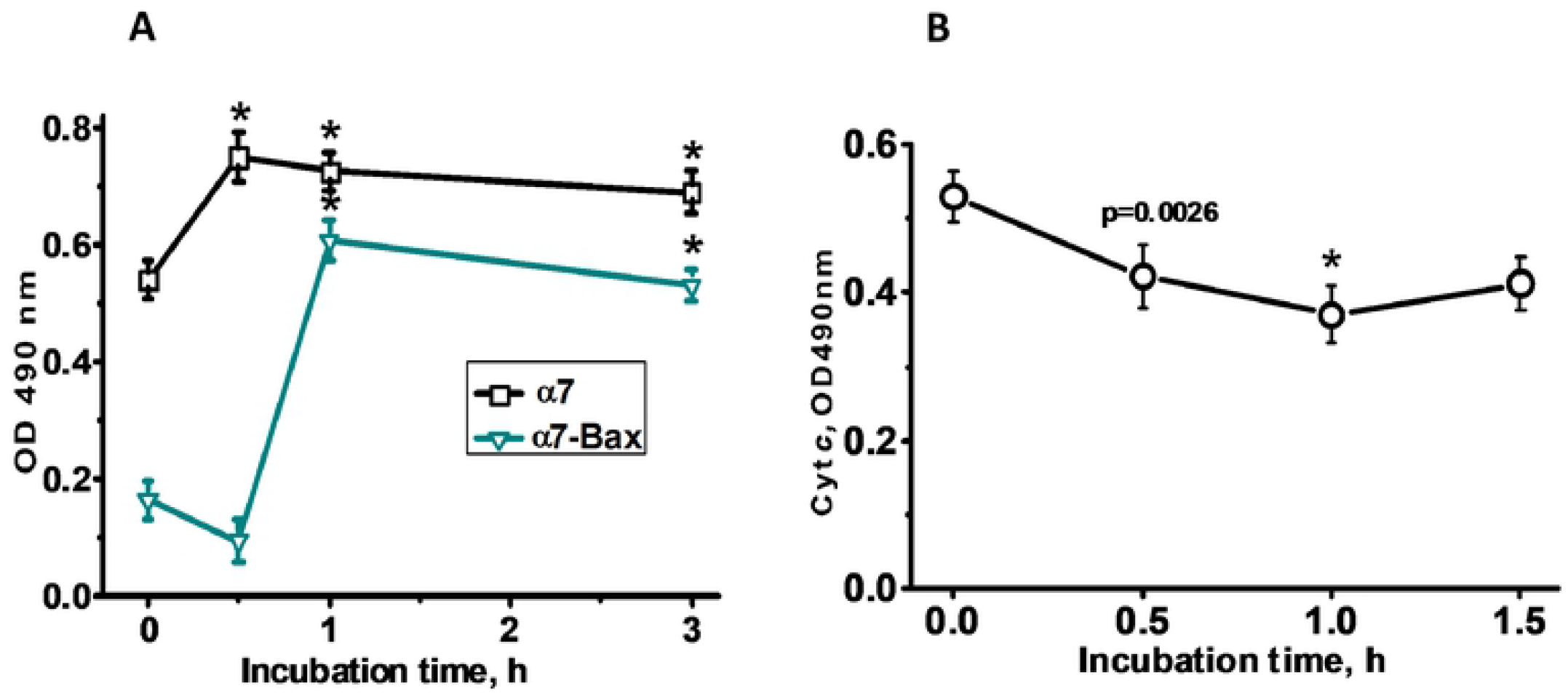
The level of α7 nAChR and α7-Bax complexes (A) and of cyt c (B) in mitochondria of U373 cells cultured in the presence of 1μM H_2_O_2_ for various periods of time. Shown are representative sandwich ELISA graphs from 3 similar experiments. * – p<0.001 compared to non-treated cells, n=3.

Our previous studies showed that both nAChR agonists and type 2 positive allosteric modulators (PAMs) attenuate cyt c release form isolated mitochondria [27]. Here we added either PNU282987 (α7-selective orthosteric agonist) or PNU120595 (α7-selective type 2 PAM) to U373 cells cultured in the presence of H_2_O_2_ for 1h and then separated the cells into mitochondria and cytosole. As shown in Fig. 4A, culturing cells in the presence of H_2_O_2_ decreased cyt c level in mitochondria and increased it in the cytosole demonstrating that cyt c was released from mitochondria into the cytosole. Addition of either PNU282987 or PNU120596 attenuated cyt c release, similarly to what we have previously shown for isolated mitochondria treated with Ca^2+^ or H_2_O_2_ [27]. To study if α7-specific ligands influence α7-Bax complexes, we isolated mitochondria from U373 cells cultured in the presence of H_2_O_2_ for 1h and treated them with either PNU282987 or PNU120596. As a positive control, mitochondria were treated with 1% SDS, which was expected to disrupt any non-covalently bound protein complexes. As shown in Fig.4B, the signals for both α7-Bax and β2-Bax were found in mitochondria of H_2_O_2_-treated U373 cells and were significantly decreased in SDS-treated mitochondria. Similar, although not so strong decrease was observed in mitochondria treated with either PNU282987 or PNU120596. Therefore, either the agonist or PAM disrupted α7-Bax and β2-Bax complexes in mitochondria of U373 cells. Finally, we compared the levels of α7 nAChR, α7-Bax and α7-VDAC in mitochondria of U373 cells cultured in the presence of H_2_O_2_ and either treated or not with PNU282987. As shown in Fig.4C, treating U373 cells with H_2_O_2_ resulted in the increase of both α7 and α7-Bax and in the decrease of α7-VDAC. Treating isolated mitochondria with PNU282987 decreased α7 and α7-Bax and increased α7-VDAC. Taken together, these data suggested that positive effect of either agonist or PAM is due to disruption of α7-Bax complexes and restoration of α7-VDAC complexes.

**Fig 4.**
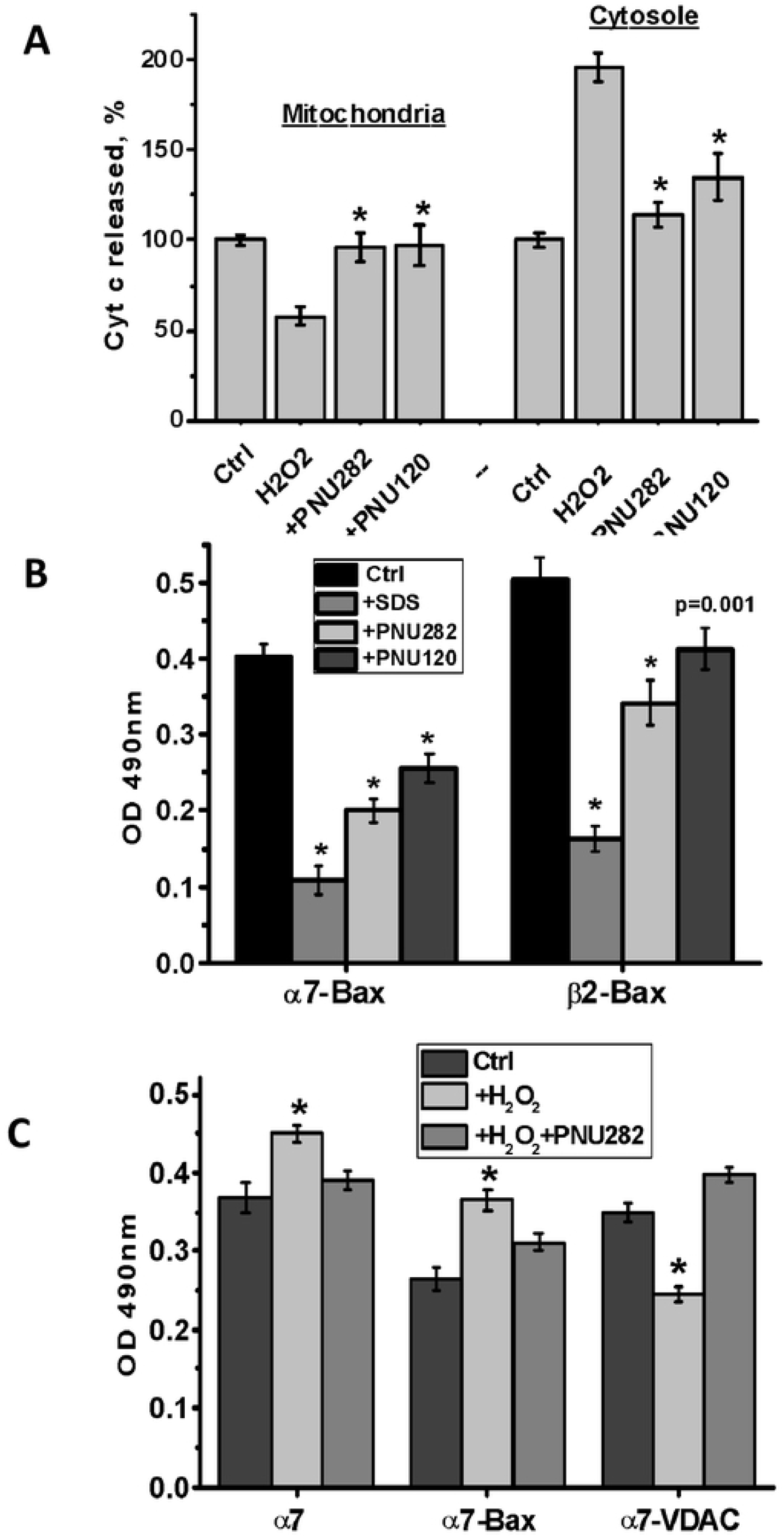
The effects of PNU282987 (PNU282) and PNU120596 (PNU120) on the level of cyt c released (A), α7-Bax and β2-Bax content (B) and the effect of PNU282 on α7-Bax and α7-VDAC (C) in mitochondria of U373 cells. **A**: The cells were pre-incubated with either PNU282 or PNU120 for 30 min, washed and treated with H_2_O_2_ for 1 h; the isolated mitochondria and remaining cytosole were analyzed by sandwich ELISA; * – p<0.001 compared to H_2_O_2_ –only treatment (4 experiments); **B**: Mitochondria isolated from U373 cells cultured with H_2_O_2_ for 1h were treated with 1% SDS, 30 nM PNU282987 or 300 nM PNU120596 for 15 min, then lyzed and studied by sandwich ELISA; * – p<0.001 or p=0.0011 compared to non-treated mitochondria; **C**: Mitochondria were isolated form U373 cells cultured with or without H_2_O_2_ for 1 h; a half of mitochondria from H_2_O_2_-treated cells was additionally treated with PNU282 for 15 min; * – p<0.001 compared to mitochondria of non-treated cells (Ctrl), n=3.

Encouraged by this data, we asked if PNU282987 can prevent apoptosis of U373 cells caused by H_2_O_2_. To our surprise, addition of PNU282987 did not improve the survival of U373 cells according to MTT test (Fig.5A). Further experiments demonstrated that PNU282987 stimulated IL-6 production in U373 cells treated with H_2_O_2_ (Fig. 5B). Addition of recombinant IL-6 did not influence U373 cells in the absence of H_2_O_2_ but obviously aggravated H_2_O_2_ effect (Fig.5C). This data allowed us to explain the absence of anti-apoptotic effect of PNU282987 in cultured U373 cells by increased IL-6 production.

**Fig 5.**
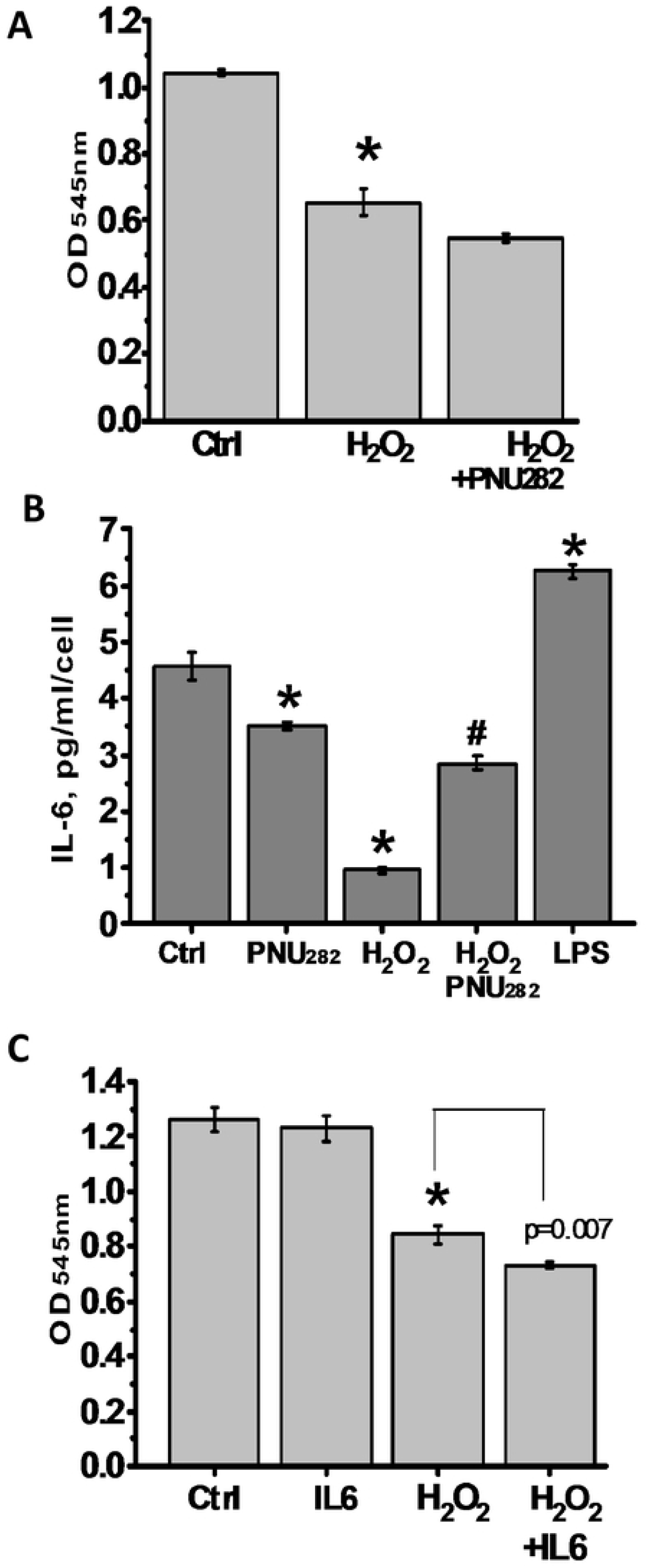
The effects of PNU282987 (PNU282) (A) or recombinant IL-6 (B) on the viability and the effect of PNU282987 on IL-6 production in U373 cells. (**C**). The cells were cultured in the presence of H_2_O_2_, PNU282, IL-6, their combinations or LPS, as described in methods. The supernatants were collected and tested for IL-6 content, while the cells viability was studied in MTT test; * – p<0.001 compared to non-treated cells (Ctrl); # – p<0.001 compared to H_2_O_2_-treated cells (n=6).

## Discussion

The data from Grando’s laboratory published in 2003 demonstrated that keratinocytes of α7 nAChR knockout mice possessed decreased amounts of the pro-apoptotic Bad and Bax at both the mRNA and the protein levels, suggesting that α7 nAChR is coupled to stimulation of keratinocyte apoptosis [28]. Our data for the first time show the direct interaction of α7 nAChRs with Bax and suggest its role in mitochondria-driven apoptosis.

In silico data shown here demonstrated that nAChRs are capable to interact with VDAC and Bax. For the modeling, we used a protein sequence of nAChR from *Torpedo marmorata.* However, high homology of nAChR subunits in their transmembrane portions [1] allows suggesting that the model is true for other nAChR subtypes, including those found in mitochondria. In most of sandwich assays presented here we used the capture antibody generated against the whole extracellular domain (1-208) of α7 subunit, which can bind almost all homologous nAChR subunits. The additional use of α7- and β2-specific antibodies showed the presence of α7-Bax and β2-Bax complexes suggesting the involvement of α7β2 nAChRs previously found in mitochondria [29]. The effects of α7-specific ligands also indicate the involvement of α7-containing nAChRs. However, ligands selective for other than α7β2 nAChR subtypes also prevented cyt c release from mitochondria [29–30]; therefore, one can expect that other mitochondrial nAChRs function similarly to α7-containing ones.

The nAChR-Bax complexes were formed within 1h after apoptosis induction and were disrupted by α7-specific agonist or type 2 PAM resulting in attenuation of cyt c release from mitochondria. The nAChR-Bax complexes formation was accompanied by disruption of nAChR-VDAC complexes, which were restored upon agonist addition. Taken together, these data demonstrate an important role of nAChRs in regulating the interplay of mitochondrial proteins involved in mitochondrial apoptosis-induced channel (MAC) formation.

Molecular processes underlying the formation of MAC [31], as well as its protein components, are still under debate. The early models suggested that pro-apoptotic factors are released from mitochondria through permeability transition pore composed of the VDAC1 located within the outer mitochondria membrane, adenine nucleotide translocase located within the mitochondrial inner membrane, and cyclophilin D located within the mitochondrial matrix [32]. Opening the pore was suggested to occur after activated Bax binds to VDAC1 [33]. However, later studies using mitochondria of VDAC-knockout animals put in doubt the role of VDAC in this process [11]. In addition, the permeability pore diameter was considered to be too small for cyt c to be released. An alternative model suggests that MAC is formed in the mitochondria outer membrane by activated Bax/Bak, which oligomerize yielding a pore diameter of 5 nm, which is sufficient to release cyt c [34]. Any of these models assumes that Bax undergo substantial conformational changes upon activation, binding to the membrane or oligomerization. Dynamic conformational changes are a feature of all three Bcl-2 families [13].

The nAChRs are also conformationally labile proteins. The pentameric structure allows significant conformational movements resulting in the ion channel opening upon agonist binding. Accompanying changes in the cytoplasmic part of nAChR subunits engage multiple kinases mediating metabotropic signaling. Previously we reported that metabotropic signaling of mitochondrial nAChRs may be stimulated by orthosteric agonists, PAMs or even antagonists and is largely dependent on the conformational changes in the nAChR molecule caused by their binding [27, 30]. Here we show one of the mechanisms through which mitochondrial nAChRs can influence the MAC formation. According to presented results, α7 nAChRs are initially bound to VDAC1 that is in accord with our previous data [8]. Apoptogenic stimulus (H_2_O_2_) increases the α7 nAChR content in mitochondria followed by α7-Bax complexes formation. It is not clear if nAChRs are translocated to mitochondria from the cytosole or are released from complexes with VDAC, which are disrupted. Our previous data demonstrated that the liver damage by partial hepatectomy results in initial decrease of nAChRs in the cytosole and their increase in mitochondria of liver cells suggesting that the nAChRs were re-distributed between the cytosole and mitochondria [35]. Therefore, it looks probable that the nAChRs arrive to mitochondria from the cytosole to bind with Bax. The time of α7-Bax complexes formation coincides with the maximum of cyt c release, indicating that α7 to Bax binding favors the MAC formation. Either orthosteric agonist (PNU282987) or type 2 PAM (PNU120596) disrupted α7-Bax complexes and attenuated cyt c release from mitochondria. The molecular modeling shows that both Bax and VDAC1 bind to transmembrane M4 α-helix of nAChR subunit. According to a structural nAChR model, the M4 portions are located on the periphery of each subunit and are exposed to the lipid bilayer [36]. Moreover, the M4 portions were suggested to be a target for the intramembrane lipid regulation profoundly affecting the nAChR functional conformations [37]. These data help understanding how the membrane-bound Bax or VDAC interact with nAChR within the mitochondria outer membrane and why PNU282987 or PNU120596 affect this interaction. PNU282987 (agonist) binds to extracellular acetylcholine-binding site resulting in gating accompanied by substantial conformational perturbations of the nAChR molecule [38–39]. PNU120596 (type 2 PAM) binds within an intrasubunit transmembrane cavity formed by M1-M4 residues [40–42] and obviously can affect M4 conformation and even replace Bax. Similar binding area of Bax and VDAC within the nAChR molecule allows suggesting that these two proteins compete for binding to α7 nAChR: the apoptotic stimulus shifts the equilibrium in favor of Bax while ligating α7 nAChR with the agonist or PAM disrupts α7-Bax and returns α7 to VDAC. Assuming that activated Bax binds to VDAC1 to open the pore [33], we can hypothesize that α7 nAChR protects VDAC from Bax binding. The details of these interactions obviously need further examination.

Finally, our data demonstrate that at the whole-cell level the effects of mitochondrial and plasma membrane nAChRs may be different.PNU292987 prevented cyt c release from mitochondria under the effect of H_2_O_2_, but stimulated IL-6 production by U373 cells that aggravated H_2_O_2_ effect.

The data obtained naturally put a question if nAChRs also interact with other pro- and anti-apoptotic Bcl-2 family proteins. Taking into account the structural homology within the family, one can expect that all of them are potentially able to bind to nAChR transmembrane portions; we have preliminary data on the existence of nAChR-Bcl-XL and nAChR-Bcl-XS complexes. Their interplay in the course of apoptosis induction is a fascinating problem to be investigated in future experiments.

## Conclusions

1. Mitochondrial nAChRs form complexes with both VDAC and Bax.
2. Apoptogenic stimulus results in translocation of α7 nAChRs to mitochondria, formation of α7-Bax complexes but disruption of α7-VDAC complexes.
3. α7-selective orthosteric agonist and type 2 PAM disrupt α7-Bax, return α7 to complex with VDAC and prevent cyt *c* release from mitochondria.
4. The data obtained suggest that mitochondrial nAChRs regulate the interplay of mitochondrial proteins involved in MAC formation.
5. The effect of mitochondrial and plasma membrane nAChRs may be different at the whole cell level.

